# Automated prediction of the clinical impact of structural copy number variations

**DOI:** 10.1101/2020.07.30.228601

**Authors:** Michaela Gaziova, Ondrej Pös, Werner Krampl, Zuzana Kubiritova, Marcel Kucharik, Jan Radvanszky, Jaroslav Budis, Tomas Szemes

## Abstract

Copy number variants (CNVs) play important roles in many biological processes, including the development of genetic diseases, making them attractive targets for genetic analysis. This led to the demand for interpretation tools that would relieve researchers, laboratory diagnosticians, genetic counselors and clinical geneticists from the laborious process of annotation and classification of CNVs. Here we demonstrate that the prediction of the clinical impact of CNVs can be automated using modern machine learning methods applied to publicly available genomic annotations, requiring only basic input information about the genomic location and structural type (duplication/deletion) of the analyzed CNV. The presented approach achieved 0.95 prediction accuracy on deletions and 0.96 on duplications from the ClinVar dataset and therefore have a great potential to guide users to more precise conclusions.

## Introduction

Copy number variants (CNVs) are unbalanced structural rearrangements of the genome which lead to variability, i.e. relative difference in the number of copies of particular DNA sequences when comparing genomes of different individuals belonging to the same or to distinct populations of a biological species (Pös et al. 2020, article in preparation…). Recent studies typically defined the size of CNVs to be >50 bp [1]. This type of variation includes deletions and duplications that can result in copy number gain or copy number loss [2]. These structural variants seem to be significant in the context of functionality and evolution [3]. Based on recent research, they contribute to population diversity and to the development of human genetic diseases [4, 5]. It is known that CNVs can directly affect the gene coding sequence and cause disruption of gene or alter gene dosage leading to development of several disorders [1]. It was also shown that CNVs can affect gene expression indirectly. Such variants have the potential to disrupt spatial organization of the genome, by altering chromatin interaction domains [6, 7]. One of the most important mediators of this phenomenon are topologically associated domains [8]. Another molecular mechanism by which rearrangements of the genome may cause disease phenotype is the unmasking of recessive mutations or functional polymorphisms when a deletion occurs [9]. Various methods have been developed for the analysis of CNVs, from conventional cytogenetic methods, through microarrays to the next generation sequencing (NGS) [10]. In recent years, NGS has become a valuable tool for clinical diagnostics and represents a sensitive and accurate approach for the detection of CNVs with a wide range of sizes. Currently, with the decreasing price of sequencing, there is a growing number of tests related to CNV detection [11]. CNV analysis is increasingly used in clinical testing through genome, exome, or gene panel sequencing. These methods have enabled genome-wide detection of CNVs in clinically affected individuals, as well as in the general population [12, 13]. Due to a significant progress in the detection of structural variants, we are now able to detect thousands of structural variants with a deep coverage sequencing in a human genome. However, since the speed of novel variants identification is greater than the speed at which interpretation tools are developed, there is a growing gap in our understanding of the clinical implications of DNA variants [12]. Moreover, until recently, detailed guidelines for CNV interpretation were not available [14, 15]. Therefore, the prediction of the functional impact and clinical relevance of such variants still remains a challenge. In the past, prediction of the impact of single nucleotide polymorphisms on the protein function met a similar problem and great effort led to the development of many tools for pathogenicity prediction [16]. Today, some of these tools can calculate a score of pathogenicity for variants located in various positions throughout the genome. However, the development of such tools for structural variants seems to be more difficult. This is because CNVs have a wide spectrum of lengths, ranging from 50 bp to several Mbp.

Moreover, the genomic coordinates highly differ, affecting various genes, regulatory or other functionally important regions. All these factors should be considered when developing a method for prediction of the impact of structural variants for appropriate prioritization and classification of such variants [17].

Tools for CNV classification and/or annotation have recently been reviewed (Pös et al. 2020 and references therein, article in preparation), although none of them evaluate the final probability of pathogenicity. Therefore, it is not possible to compare the accuracy of prediction between tools [17].

In the present study, we demonstrate that the prediction of the clinical impact of structural variation can be automated to relieve researchers, laboratory diagnosticians, genetic counselors and clinical geneticists from the tedious interpretation process.

## Materials and Methods

CNVs downloaded from the Clinvar database [18] are annotated with relevant attributes gathered from several publicly available databases. These attributes are then aggregated and transformed into a feature vector. Finally, the pre-trained classifier scores the vector with the probability of CNV pathogenicity, with 0% meaning no probability of pathogenicity and 100% meaning high suspicion of pathogenicity of a particular CNV. Main steps of this prediction process, together with our testing approach, are described in the following sections.

### Annotation

At first, query CNV is annotated with attributes gathered from several databases (see chapter Selection of attributes). Currently, coding regions of the genome represent the most prominent source of relevant attributes, although other data sources, e.g. variants, promotors, known CNV cases etc. may be utilized in the same way. Genes covered by the query CNV are extracted. All genes from the set are then annotated with additional attributes from publicly available databases to describe their isolated effect on the phenotype. Gene attributes are then aggregated to identify extremal values. Each attribute represents a different view on the affected genomic region and the potential cause of genetic disease. All gathered attributes of the CNV are joined into a numeric feature vector and passed to the classification step of the analysis (Fig.1).

**Figure 1.**
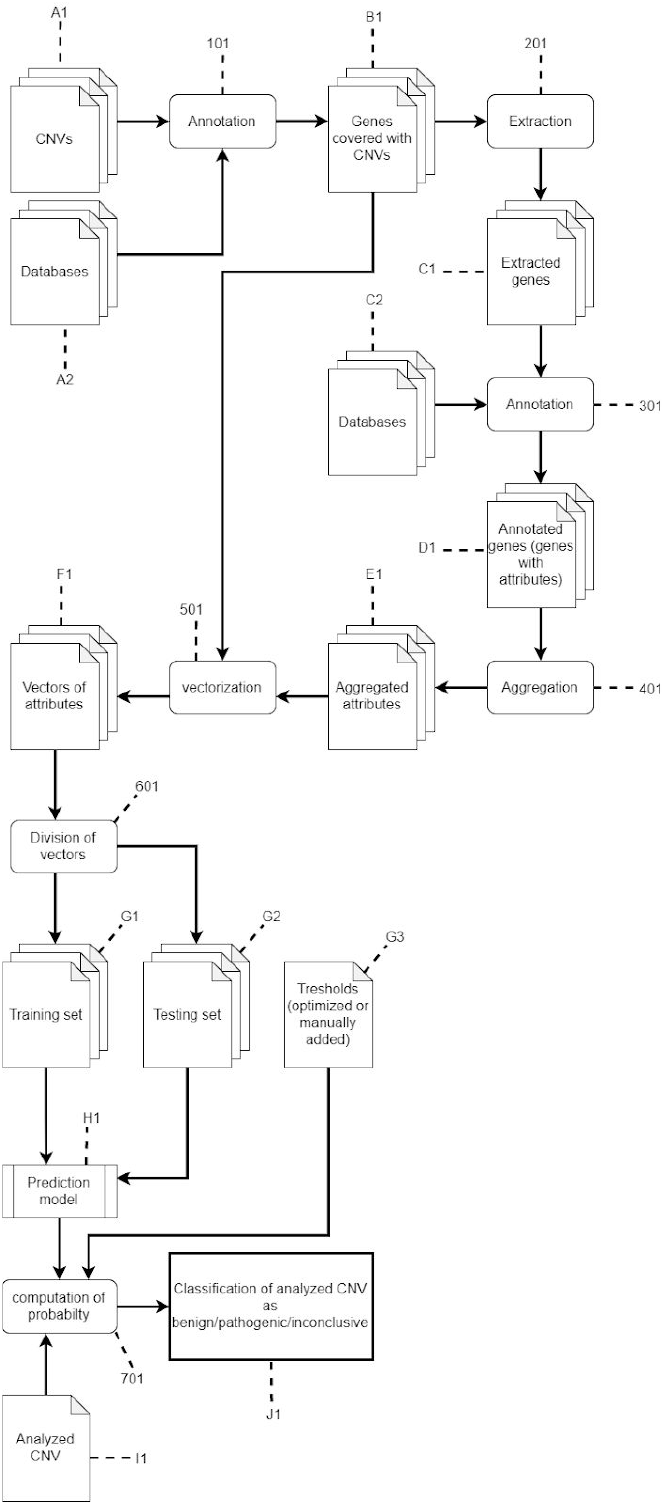
Detailed scheme of the algorithm for the CNV classification method. CNV (A1) is annotated (101) with attributes from several databases (A2). Genes (C1) covered by the CNV (B1) are extracted (201) and annotated (301) with additional attributes from public databases (C2) to describe their effect on the phenotype. Gene attributes (D1) are aggregated (401), to identify extremal values (E1). All gathered attributes of the CNV are joined (501) into a numeric feature vector (F1) and passed to the classification step of the analysis. Features vectors are randomly divided (601) into two disjunct sets: the training set (G1) for the learning of the classifier, and the testing set (G2) for evaluation of its accuracy. New CNV, which is not part of the Training or Testing set, called Analyzed CNV (I1) is then analyzed (701) on the trained model (H1) and a probability and classification (benign/pathogenic/inconclusive) are yielded (J1).

### Selection of attributes

Data for training and validation of the method’s prediction model were acquired from the ClinVar database [18] that represents a shared source of observed genomic variation collected from various studies and individual patients. Only structural deletions (27,411 CNVs) and duplications (26,839 CNVs) were extracted with clearly defined clinical significance: either a pathogenic or a benign (18,149 duplications and 12,509 deletions). Clinical significance of each CNV was provided by ClinVar curation. Part of the data was removed due to bad format and wrong values of CNV coordinates. As a subsequent step, structural variations were annotated and ranked with the AnnotSV tool [19]. CNVs as well as each overlapped gene were annotated with 72 attributes gathered mostly from population studies and genomic databases such as DDD (Deciphering Developmental Disorders) [20], IMH (Ira M. Hall’s lab) [21], 1000 Genomes project [22], OMIM (Online Mendelian Inheritance in Man) [23], ExAC (Exome Aggregation Consortium) [24], and DGV (Database of Genomic Variants) [25].

Dataset was further divided into pathogenic (6,011) and benign (12,004) deletions, and pathogenic (2,872) and benign (9,391) duplications to obtain information on how pathogenic and benign CNVs differ from each other. Each numerical attribute was examined to identify those that significantly differ between the groups, and thus may be instructive in the classification. Since attributes were typically not normally distributed, nonparametric Mann-Whitney U test was used to evaluate their significance. For further analyses, only attributes with a p-value lower than 0.001 were chosen. Attributes with missing values were also filtered. Those attributes that played an important role in discriminating between CNV pathogenicity and benignity were selected (Table 1) Differences between pathogenic and benign CNVs in attributes are shown in Supplementary Fig. 1.

**Table 1.** The list of attributes used for CNV annotation and training of the prediction model. It is possible to extend the list with other predictive attributes to improve overall model accuracy.

| Attribute | Definition | Prediction model training |  |
| --- | --- | --- | --- |
|  |  | Del | Dup |
| Gene count | The number of covered genes | Yes | Yes |
| Morbid gene count | The number of genes that are intolerant to an irregular number of copies according to the OMIM database [23] | Yes | Yes |
| EpLI score | Number of covered genes with high probability ($pLI \geq 0.9$) that a gene is intolerant to a loss of function mutation. Genes with $pLI \geq 0.9$ are extremely intolerant to loss of function mutation according to ExAC study [24]. The pLI score represents intolerance of a gene to a loss of function mutation | Yes | Yes |
| HI score | Counts of overlapped genes with low haploinsufficiency score ($HI \leq 10$). The lower value represents the higher intolerance of a gene to retain normal function in a case of its deletion. HI scores of genes are provided by the DDD study [20] | Yes | No |
| CG score | Counts of genes with high haploinsufficiency (triplosensitivity) score (2 or 3) for deletion (duplication) event provided by ClinGen consortium rating system [27] | Yes | No |
| DGV Frequency | Minimal frequency of genes in individuals sequenced in the DGV study [25]. Lower values represent less conserved genes with impact on an irregular copy number | Yes | Yes |
| EZ score | Maximal Z score of genes overlapped by CNV. Higher Z score means higher gene intolerance to loss of function mutation of CNVs. Z scores are provided by ExAC study [24] | Yes | Yes |

### Classification

Given a feature vector produced in the annotation step, a pre-trained classifier calculates a probability of the CNV pathogenicity. The predicted condition (benign/pathogenic) may be determined from the probability in several ways. A CNV is standardly considered as pathogenic when the calculated probability is above 50%. Since most of the wrong predictions are expected for CNVs near this threshold, it is possible to choose arbitrary lower (e.g. 20%) and upper (e.g. 80%) thresholds to eliminate uncertain predictions. CNVs with probabilities that fall between those thresholds are considered as inconclusive. The thresholds may be also estimated automatically, using additional optimization steps following the training step. If there is an acceptable proportion of inconclusive samples (e.g. 5%), the method then finds such thresholds that will eliminate at most 5% samples, while maximizing the overall accuracy of the rest of the samples. Alternatively, required accuracy (e.g. 0.98) for identification of optimal thresholds can be supplied manually.

### Training and testing

The classification method was trained on a set of CNVs with known clinical impact. Such a set is publicly available from the ClinVar database [18]. CNVs of different structural types (e.g. deletion, duplication) should be trained and tested separately, resulting in dedicated classifiers for each structural type. Subsequently, the dataset for the training of the method was divided into two disjunct sets. Training set (80%, consisting of 4,847 pathogenic and 9,565 benign deletions and 2,288 pathogenic and 7,522 benign duplications) for the learning of the classifier and testing set (20%, consisting of 1,164 pathogenic and 2,439 benign deletions, 584 pathogenic and 1,869 benign duplications) for evaluation of its accuracy on an independent set of CNVs. Attributes for the method were chosen through recursive elimination and according to significance of the features in the decision function (Supplementary Table 1a, 1b). Several classification algorithms (such as XGBoost Classifier, Support Vector Classifier, Random Forest Classifier, and others) based on the training set were tested. Following the accuracy of the predictor on the testing set (summary of metrics can be seen in Supplementary Table 2a, 2b, Fig. 2a, 2b), a few of the tested prediction models were chosen and tested with different parameters of decision function. Various parameter sets were examined, also with the help of the ROC curve and Grid Search (tuning technique of hyperparameters) (Supplementary Fig. 8). Finally, a random forest predictor algorithm implemented in XGBoost Python3 module [26] was chosen as the classifier of pathogenicity of CNVs. Parameters for training the model were set to default values except for the learning rate (0.5 for deletions and 0.6 for duplications). Dataset for training and testing is available at github: https://github.com/MiskaGaziova/ISV.

**Figure 2.**
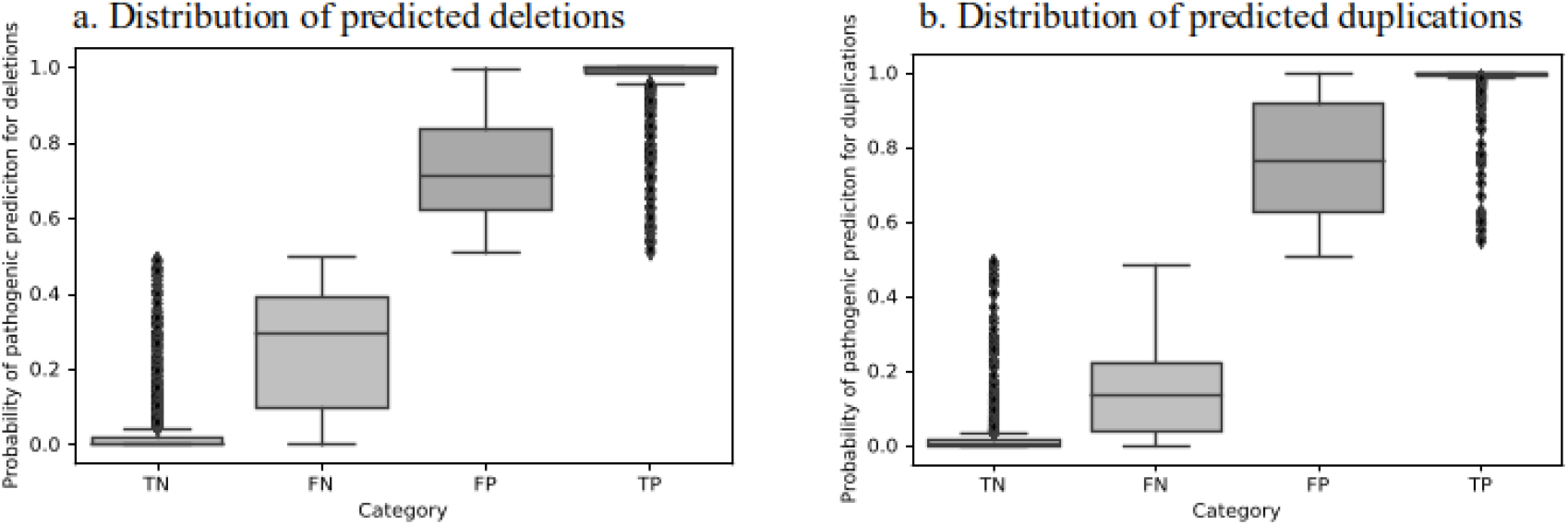
Distribution of predicted CNVs on testing data (Category) according to its probability of pathogenic prediction. (TN = True negative, FN = False negative, TP = True positive, FP = False positive) (Distribution of predicted CNVs on training data can be found in Supplementary Fig. 3)

### Identification of optimal thresholds for inconclusive predictions

Implemented classifier returns a probability of pathogenic clinical impact of the examined CNV. The samples with probability around 50% are more suspicious of wrong predictions, and therefore should be concluded as inconclusive. We employed the following strategy to identify optimal thresholds for the elimination of at most 5% samples that maximize overall accuracy:

Firstly, the processed data was sorted in order to the evaluated probability of pathogenicity. Thus, we obtained an ordered sequence of CNVs from benign to pathogenic prediction. Secondly, the incorrectly predicted CNVs were counted in every 5% interval by shifting that 5-percent interval (interval represents 180 deletions and 122 duplications from the testing set) (Supplementary Fig. 4). Next, new datasets were made by removing the aforementioned interval. Then, we evaluated how the accuracy, precision, sensitivity, and specificity of the classifier changed on the new dataset after removing the 5-percent interval. The accuracy of the prediction was highest when the interval of deletions with pathogenic predictions around 50% is removed as well as the interval of duplications with pathogenic predictions around 40% (Supplementary Fig. 5). Subsequently, we found probability of pathogenic predictions of removed CNVs to set borders of pathogenic prediction for classifying as ‘inconclusive’.

## Results

### Selection of attributes

Each attribute represents a different view on the affected genomic region and the potential cause of genetic disease. All gathered and aggregated attributes of the CNV are joined into a numeric feature vector and passed to the classification step of the analysis. The most important attributes, whose variables differ between pathogenic CNVs (were used for training of the prediction model) are listed in Table 1 (all attributes are in Supplementary Table 1a, 1b).

We found out that the main risk was the length of CNV as an attribute, which increased the probability of CNV pathogenicity and that resulted in a high pathogenicity score (increased count of false positive predictions). According to the importance of the features in the decision function, the best predictive attribute for deletions is ‘CG score’ and for duplications is the ‘count of genes’ (order of significance of attributes can be seen in Supplementary Table 4).

### Prediction accuracy

Following the completion of the training algorithm on the ClinVar derived training set of 14,412 deletions, the accuracy for deletions reached the score of 0.96, precision 0.96, sensitivity of 0.92, and specificity 0.98. The evaluation of the method on the independent testing set of 3,603 CNVs showed an accuracy of 0.95, precision of 0.95, sensitivity of 0.92, and specificity of 0.97 (Fig. 2a). (Other metrics can be seen in Supplementary Table 3)

We managed to do similar steps on a dataset of 12,263 duplications from ClinVar, including 9,810 duplications in the training set and 2,453 in the testing set. After training on duplications on the training dataset we have gained 0.98 accuracy, 0.99 precision, 0.92 sensitivity, and 0.99 specificity. On testing set accuracy reached 0.96, precision 0.96, sensitivity 0.90, and specificity 0.98 (Fig. 2b). (Other metrics can be seen in Supplementary Table 3)

### Effect of eliminating uncertain predictions

We decided to interpret pathogenic predictions of deletions from 31% to 72% as uncertain significance, since the 5% interval with the highest count of wrong predictions contained deletions with prediction power of pathogenicity in that range. Following this, the accuracy on the training set had risen to 0.97, precision to 0.98, sensitivity to 0.95, and specificity to 0.99. On the testing set, accuracy reached 0.97, precision 0.97, sensitivity 0.95, and specificity 0.98 (Supplementary Table 3a).

For deletions the interval 13.29% – 61.8% was classified as uncertain significance using the same criteria as with deletions. Then, the overall accuracy of the prediction model on duplications on the training set has improved to 0.99, precision 0.99, sensitivity 0.97, and specificity 0.99. On testing set accuracy achieved 0.98, precision 0.99, sensitivity 0.97, and specificity 0.99 (Other metrics can be seen in Supplementary Table 3). (Distribution of predicted CNVs according to its probability of pathogenic prediction can be seen in Supplementary Fig. 7a-d)

## Discussion

Several tools have been proposed for CNVs characterization, interpretation, or annotation. Each tool provides specific information contributing to CNVs interpretation and a better understanding of the functional impact of such variants, however, they have various limitations. Many of these tools do not provide final annotation related to clinical significance and in the clinical or research setting it is inevitable to aggregate information from multiple such tools for accurate interpretation of analyzed CNVs. Still, the biggest limitation concerns the classification of variants. Moreover, CNV prediction programs have shown high false-positive CNV counts, which is the major limiting factor for the applicability of these programs in clinical studies [28].

Therefore, we aimed to design and create an automated method encompassing various parameters based on whether it would be possible to classify CNVs in the context of clinical significance. The presented method predicts the clinical impact of CNVs that may fall into the benign or pathogenic category. The method needs only the basic information about the position and type of CNV.

We have shown that this automated approach can achieve great accuracy on deletion (0.95) and duplications (0.96). Furthermore, when we allow for an acceptable portion of inconclusive samples (e.g. 5%), the accuracy further improves to 0.97 for deletions and 0.99 for duplications. None of the tools we have met so far have shown such good accuracy. On CNVs obtained from Clinvar database (genome reference GRCh37/hg19) with clinical significance ‘benign’ or ‘pathogenic’, AnnotSV tool reaches the accuracy of 0.71. Balanced accuracy of SG-ADVISER CNV tool reaches the score of 0.90 - 0.94 on data was derived from ClinVar, Decipher and others [29]. Achieved results suggest that our proposed method has a potential to be used in research and everyday clinical care.

One of the limitations of the described method is that it is not able to predict which disease is caused by the CNV classified as pathogenic. However, users could primarily look at which of the predictive attributes had increased the probability of pathogenicity. In this way, the values of the predictive attributes can lead users to the cause of pathogenicity and thus to the most likely diagnosis. The other limitation of the method is that it predicts the probability of pathogenicity only for a CNV locating on one copy of the chromosome.

We have shown that the clinical impact of a CNV can be predicted with high accuracy using aggregated information about genes overlapping the CNV from available resources, facilitating and supporting thus the decision process of the users when evaluating results of genomic analyses. Using machine learning methods, moreover, the process can be automated further decreasing the time needed for the evaluation of CNVs.

## Supporting information

Supplemental Summary (description and other Figures&Tables)

Table 1a. Feature importances of attributes for xgboost prediction model for deletions

Table 1b. Feature importances of attributes for xgboost prediction model for duplications

Table 2a. Summary of metrics from various classifiers training of deletions

Table 2b. Summary of metrics from various classifiers training of duplications

Table 3. Metrics of classifier (described in Methods) used for predicting pathogenicity CNV

Figure 2a. Accuracy of tested predictor model on deletions and duplications

Figure 2b. Precision of tested predictor model on deletions and duplications

## Acknowledgments

All authors are employees of Geneton Ltd., where they also participate in development of a commercial application for the annotation and interpretation of CNV. The presented method was filed as a patent application under the number PCT / EP2020 / 025292. Apart from the above mentioned all authors have declared no conflicts of interest.

The presented work was supported by the the Slovak Research and Development Agency (grant ID APVV-18-0319) (20% of charges) and the “REVOGENE - Research centre for molecular genetics” project (ITMS 26240220067) supported by the Operational Programme Research and Development funded by the ERDF (80% of charges).

## References

1. Nowakowska B (2017) Clinical interpretation of copy number variants in the human genome. J Appl Genet 58:449–457

2. Feuk L, Carson AR, Scherer SW (2006) Structural variation in the human genome. Nat Rev Genet 7:85–97

3. Escaramís G, Docampo E, Rabionet R (2015) A decade of structural variants: description, history and methods to detect structural variation. Brief Funct Genomics 14:305–314

4. Sudmant PH, Mallick S, Nelson BJ, et al (2015) Global diversity, population stratification, and selection of human copy-number variation. Science 349:aab3761

5. Mikhail FM (2014) Copy number variations and human genetic disease. Curr Opin Pediatr 26:646–652

6. Lupiáñez DG, Kraft K, Heinrich V, et al (2015) Disruptions of topological chromatin domains cause pathogenic rewiring of gene-enhancer interactions. Cell 161:1012–1025

7. Dixon JR, Selvaraj S, Yue F, Kim A, Li Y, Shen Y, et al (2012) Topological domains in mammalian genomes identified by analysis of chromatin interactions. Nature 485:376–380

8. Spector JD, Wiita AP (2019) ClinTAD: a tool for copy number variant interpretation in the context of topologically associated domains. J Hum Genet 64:437–443

9. Stankiewicz P, Lupski JR (2010) Structural Variation in the Human Genome and its Role in Disease. Annual Review of Medicine 61:437–455

10. Martin CL, Kirkpatrick BE, Ledbetter DH (2015) Copy number variants, aneuploidies, and human disease. Clin Perinatol 42:227–42, vii

11. (2014) Annotating DNA Variants Is the Next Major Goal for Human Genetics. Am J Hum Genet 94:5–10

12. (2014) Annotating DNA Variants Is the Next Major Goal for Human Genetics. Am J Hum Genet 94:5–10

13. Pös O, Budis J, Kubiritova Z, Kucharik M, Duris F, Radvanszky J, et al (2019) Identification of Structural Variation from NGS-Based Non-Invasive Prenatal Testing. Int J Mol Sci. https://doi.org/10.3390/ijms20184403

14. Kearney HM, Thorland EC, Brown KK, Quintero-Rivera F, South ST, Working Group of the American College of Medical Genetics Laboratory Quality Assurance Committee (2011) American College of Medical Genetics standards and guidelines for interpretation and reporting of postnatal constitutional copy number variants. Genet Med 13:680–685

15. Riggs ER, Andersen EF, Cherry AM, et al (2019) Technical standards for the interpretation and reporting of constitutional copy-number variants: a joint consensus recommendation of the American College of Medical Genetics and Genomics (ACMG) and the Clinical Genome Resource (ClinGen). Genet Med 1–13

16. Thusberg J, Olatubosun A, Vihinen M (2011) Performance of mutation pathogenicity prediction methods on missense variants. Hum Mutat 32:358–368

17. Ganel L, Abel HJ, Hall IM (2017) SVScore: an impact prediction tool for structural variation. Bioinformatics 33:1083–1085

18. Landrum MJ, Lee JM, Benson M, et al (2018) ClinVar: improving access to variant interpretations and supporting evidence. Nucleic Acids Res 46:D1062–D1067

19. Geoffroy V, Herenger Y, Kress A, Stoetzel C, Piton A, Dollfus H, et al (2018) AnnotSV: an integrated tool for structural variations annotation. Bioinformatics 34:3572–3574

20. Firth HV, Wright CF, DDD Study (2011) The Deciphering Developmental Disorders (DDD) study. Dev Med Child Neurol 53:702–703

21. Abel HJ, Larson DE, Chiang C, et al Mapping and characterization of structural variation in 17,795 deeply sequenced human genomes. https://doi.org/10.1101/508515

22. 1000 Genomes Project Consortium, Auton A, Brooks LD, et al (2015) A global reference for human genetic variation. Nature 526:68–74

23. OMIM - Online Mendelian Inheritance in Man. https://omim.org/. Accessed 28 Jul 2020

24. ExAC browser. http://exac.broadinstitute.org. Accessed 28 Jul 2020

25. MacDonald JR, Ziman R, Yuen RKC, Feuk L, Scherer SW (2014) The Database of Genomic Variants: a curated collection of structural variation in the human genome. Nucleic Acids Res 42:D986–92

26. Chen T, Guestrin C (2016) XGBoost: A Scalable Tree Boosting System. In: Proceedings of the 22nd ACM SIGKDD International Conference on Knowledge Discovery and Data Mining. ACM, New York, NY, USA, pp 785–794

27. Rehm HL, Berg JS, Brooks LD, et al (2015) ClinGen — The Clinical Genome Resource. New England Journal of Medicine 372:2235–2242

28. Samarakoon PS, Sorte HS, Stray-Pedersen A, Rødningen OK, Rognes T, Lyle R (2016) cnvScan: a CNV screening and annotation tool to improve the clinical utility of computational CNV prediction from exome sequencing data. BMC Genomics 17:51

29. Erikson GA, Deshpande N, Kesavan BG, Torkamani A (2015) SG-ADVISER CNV: copy-number variant annotation and interpretation. Genet Med 17:714–718

