## Supplemental Summary (description and other Figures&Tables) for "Automated prediction of the clinical impact of structural copy number variations"

**Supplementary**

**Table 1a.** Feature importances of attributes for xgboost prediction model for deletions

**Table 1b.** Feature importances of attributes for xgboost prediction model for duplications

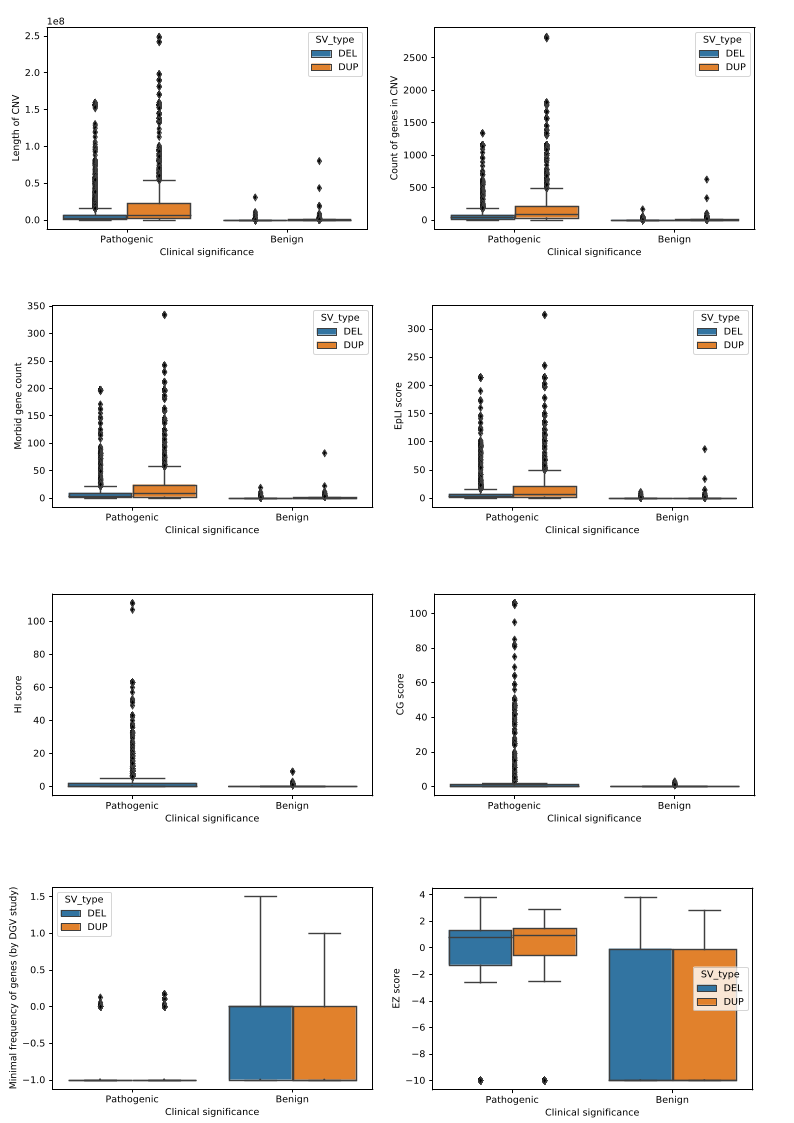

**Figure 1.** Visualisation of differences in attributes between pathogenic and benign CNVs

**Table 2a.** Summary of metrics from various classifiers training of deletions

Table shows differences in correctness of various classifiers

**Table 2b.** Summary of metrics from various classifiers training of duplications

Table shows differences in correctness between various classifiers

**Table 3.** Metrics of classifier (described in Methods) used for predicting pathogenicity CNV

**Figure 2a.** Accuracy of tested predictor model on deletions and duplications

**Figure 2b.** Precision of tested predictor model on deletions and duplications

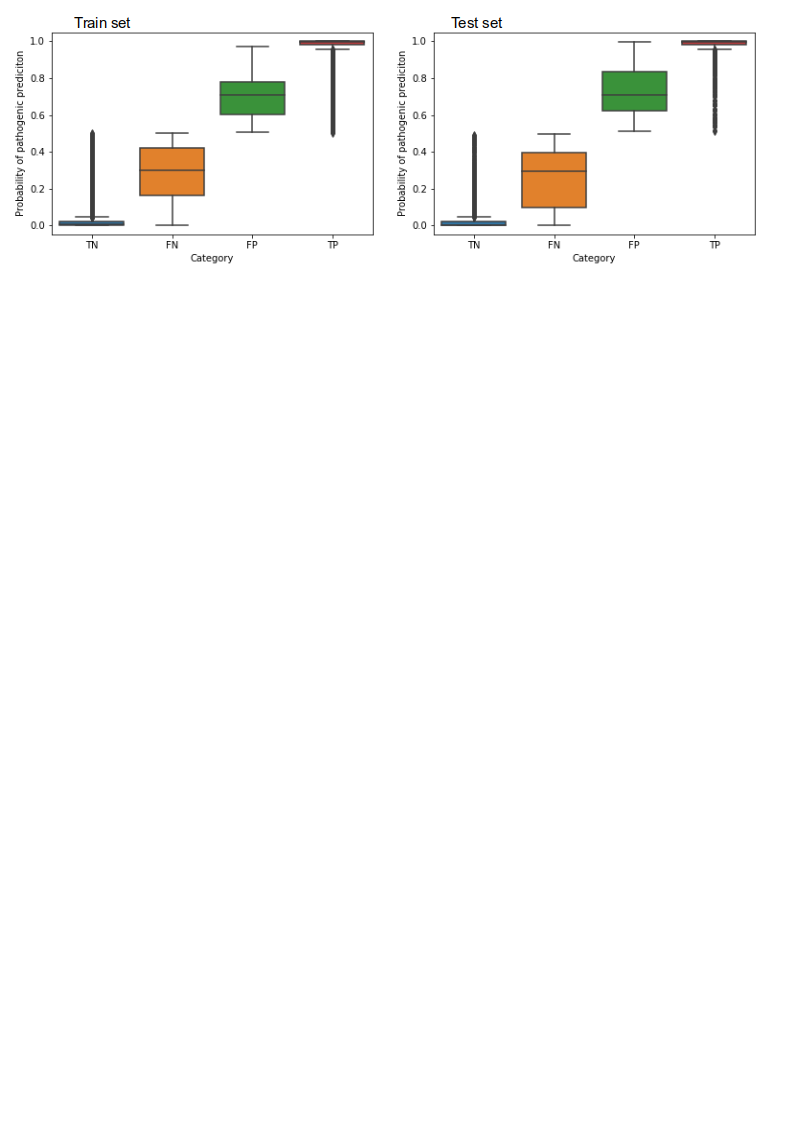

**Figure 3a**. Distribution of predicted deletions (Category) according to its probability of pathogenic prediction. (TN = True negative, FN = False negative, TP = True positive, FP = False positive)

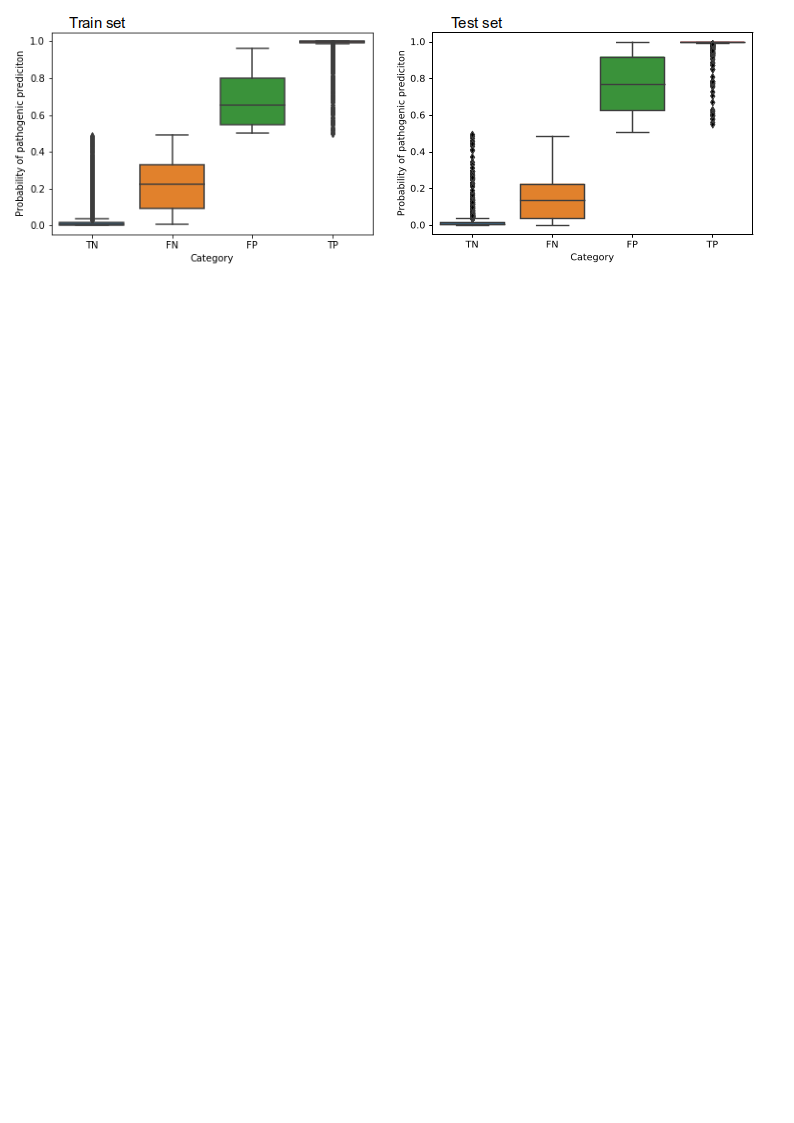

**Figure 3b.** Distribution of predicted duplication (Category) according to its probability of pathogenic prediction. (TN = True negative, FN = False negative, TP = True positive, FP = False positive)

**Identification of optimal thresholds for inconclusive predictions**

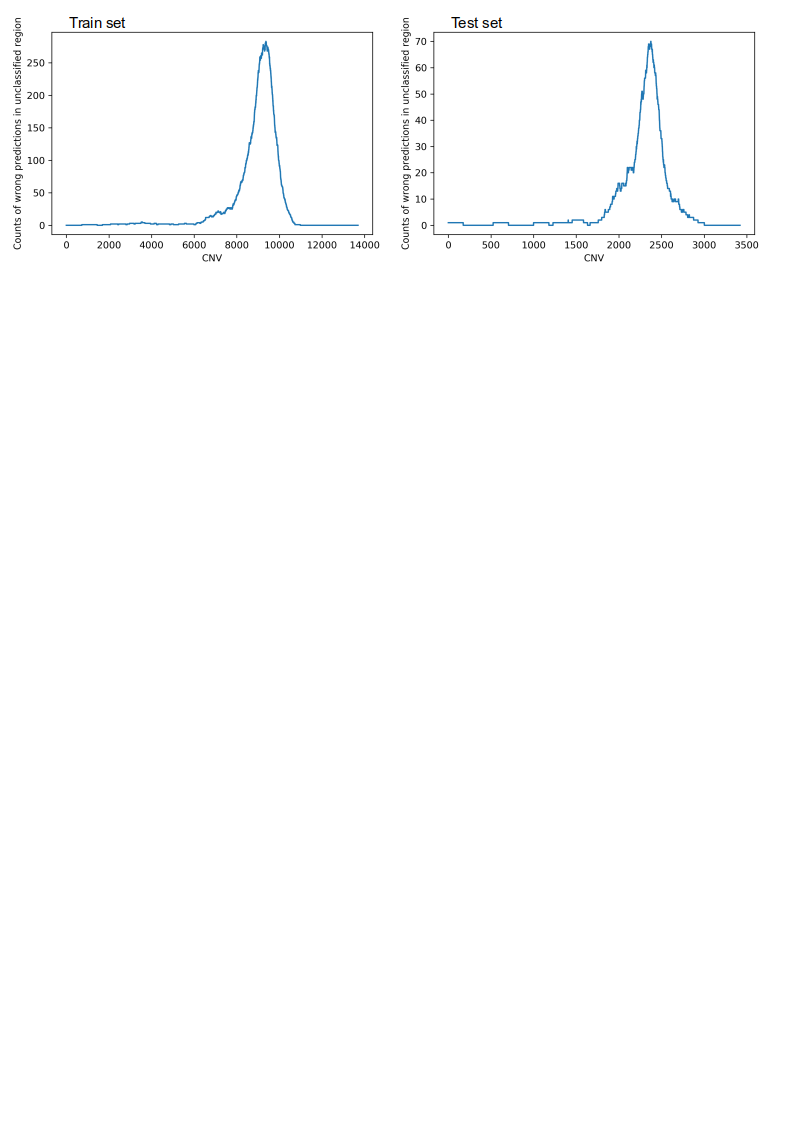

**Figure 4a.** Sorted CNV (deletions) dataset according to predicted pathogenicity and their counts of incorrectly predicted CNV in (simulated) unclassified region (CNV = index of sample)

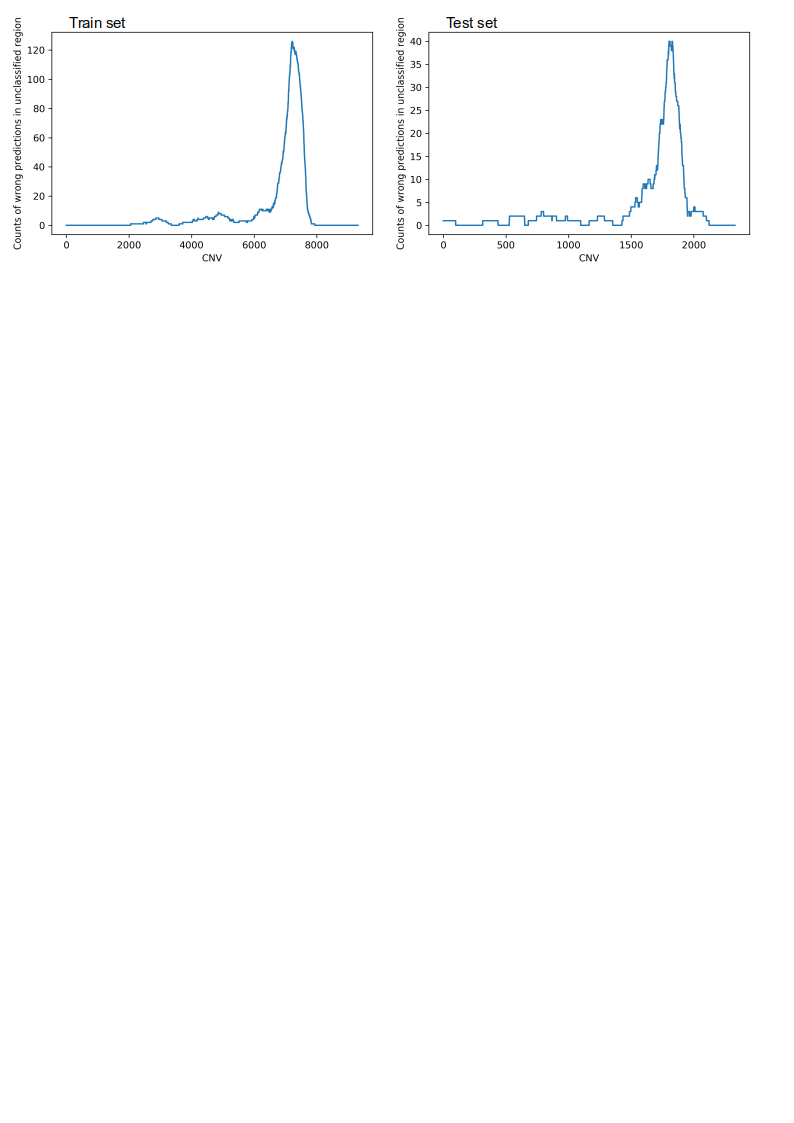

**Figure 4b.** Sorted CNV (duplications) dataset according to predicted pathogenicity and their counts of incorrectly predicted CNV in (simulated) unclassified region (CNV = index of sample)

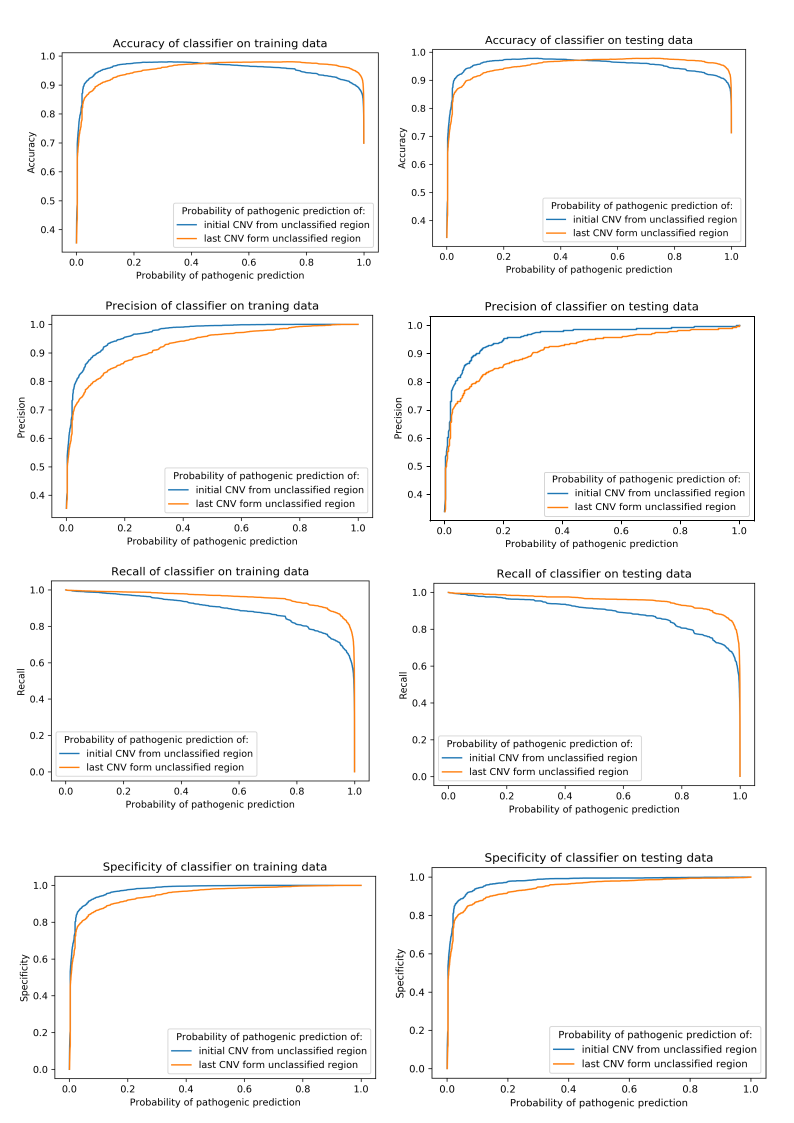

**Figure 5a.** The graphs (sest of graphs) show how the accuracy, precision,recall and specificity changes as a result of removing the 5% interval of samples from the data of deletions.

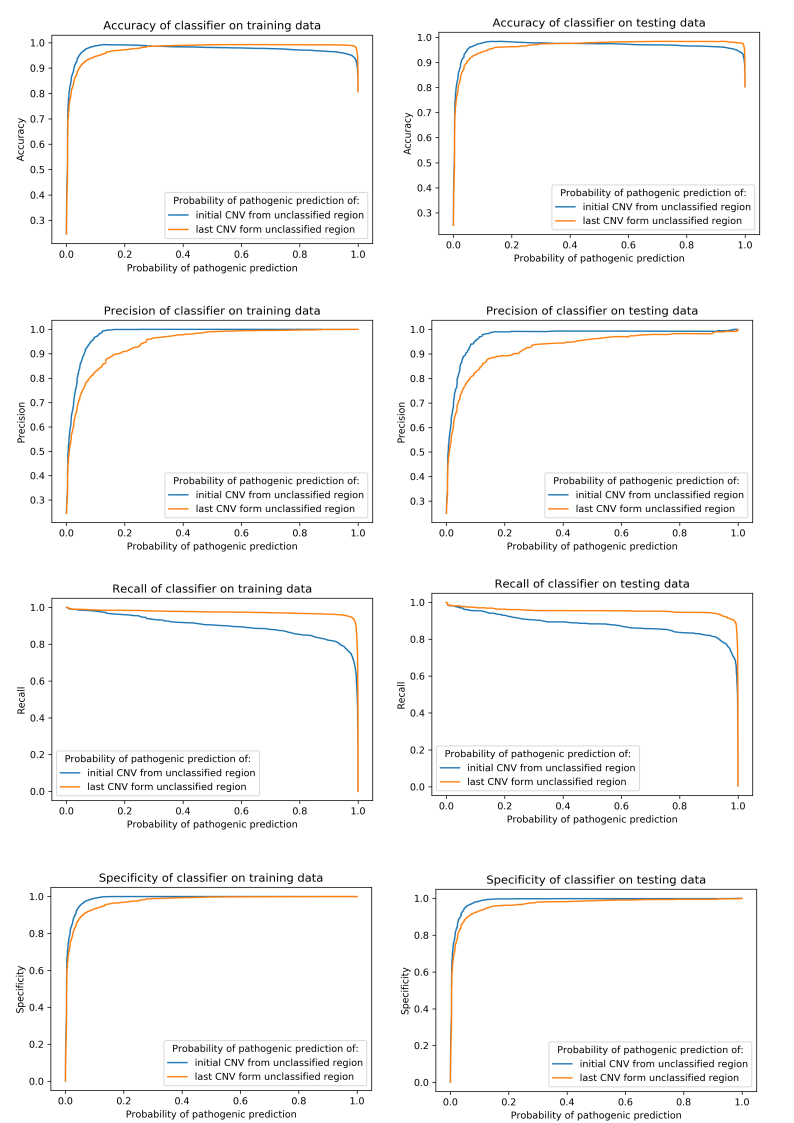

**Figure 5b.** The graphs (sest of graphs) show how the accuracy, specificity, precision and recall changes as a result of removing the 5% interval of samples from the duplications data.

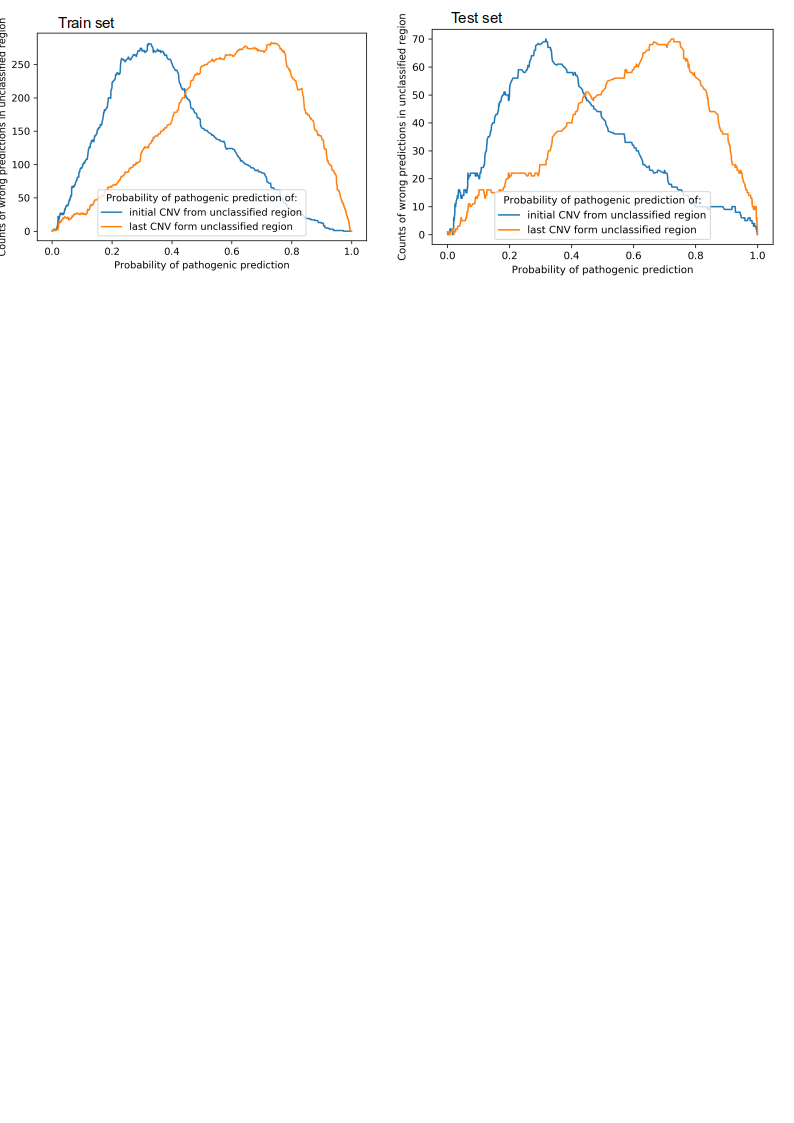

**Figure 6a.** Figure shows how pathogenicity prediction changes according to initial and last CNV sample on deletion dataset in removed 5% interval.

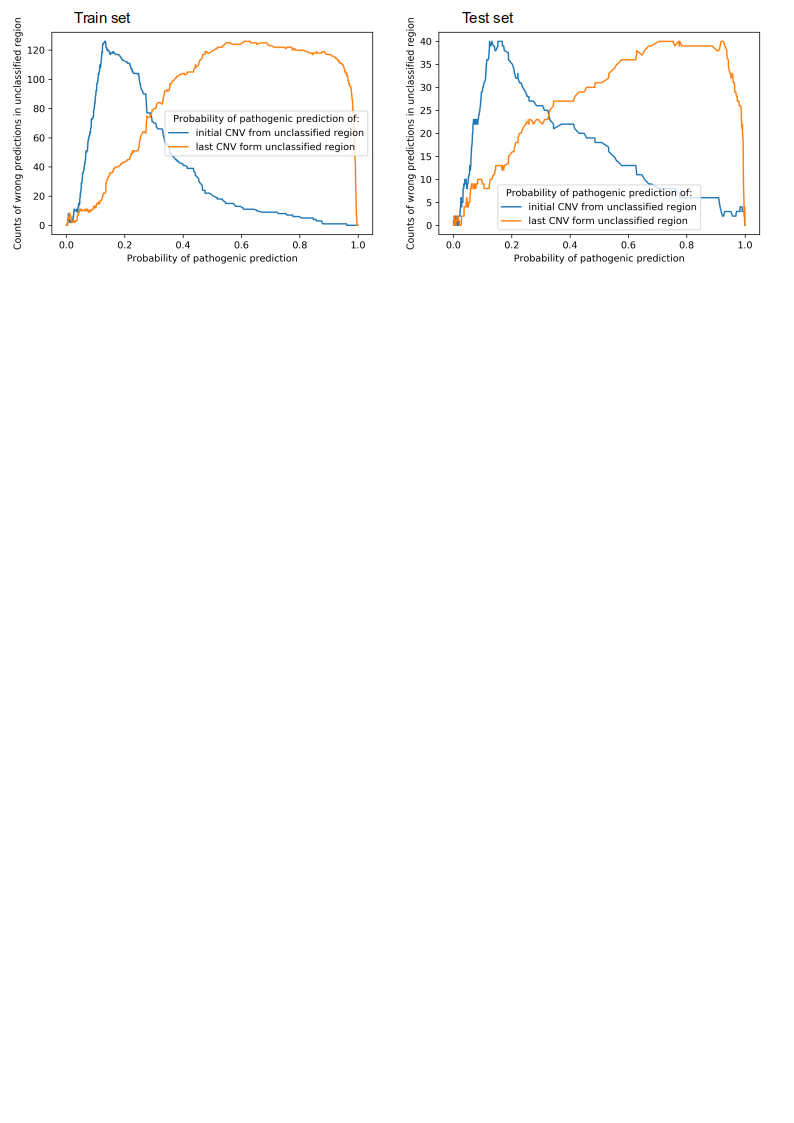

**Figure 6b.** Figure shows how pathogenicity prediction changes according to initial and last CNV sample on duplication data in removed 5% interval.

|  | **accuracy** | **precision** | **sensitivity** | **specificity** |
| --- | --- | --- | --- | --- |
| **test set** | 0.9780701754385965 | 0.976145038167939 | 0.9534016775396086 | 0.9893481039625053 |
| **train set** | 0.979947306791569 | 0.9830002297266253 | 0.9553471757088636 | 0.991943385955362 |

**Table 3a.** Summary of metrics after removing deletions with pathogenic prediction between 31-72%. That range was determined by test set. Range of pathogenic prediction on train set resulted in 32 – 73%

|  | **accuracy** | **precision** | **sensitivity** | **specificity** |
| --- | --- | --- | --- | --- |
| **Test set** | 0.9816631130063966 | 0.9701492537313433 | 0.9506398537477148 | 0.9911012235817576 |
| **Train set** | 0.9927038626609442 | 0.99476688867745 | 0.973463687150838 | 0.9984662576687117 |

**Table 3b.**

Summary of metrics after removing duplications with pathogenic prediction between 13.29-61.8% .

That range was firstly determined on test set. However, that range of pathogenic prediction evaluated on test set (13.57-77.6) exceed 5% of train set and as we can’t remove more than 5% of data, we decided to use range of pathogenic prediction according to train set, which is 0.1329 – 0.618%.

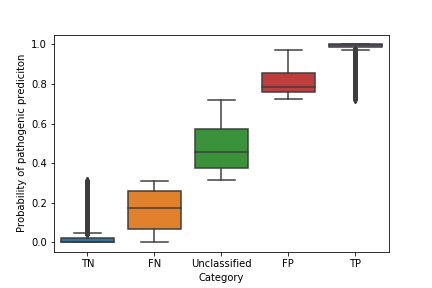

**Figure 7a.** Distribution of predicted deletion on training data (Category) according to its probability of pathogenic prediction. (TN = True negative, FN = False negative, TP = True positive, FP = False positive)

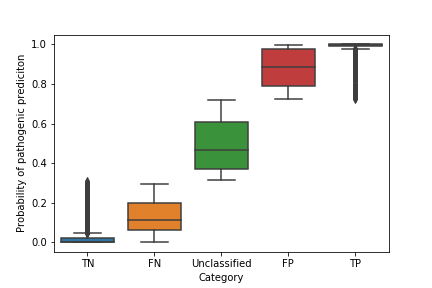

**Figure 7b.** Distribution of predicted deletion on testing data (Category) according to its probability of pathogenic prediction. (TN = True negative, FN = False negative, TP = True positive, FP = False positive)

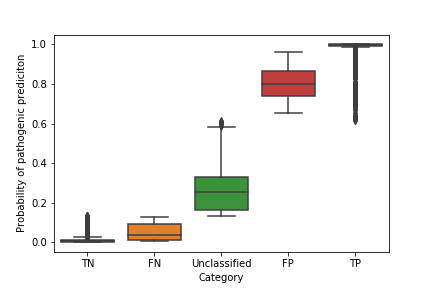

**Figure 7c.** Distribution of predicted duplication on training data (Category) according to its probability of pathogenic prediction. (TN = True negative, FN = False negative, TP = True positive, FP = False positive)

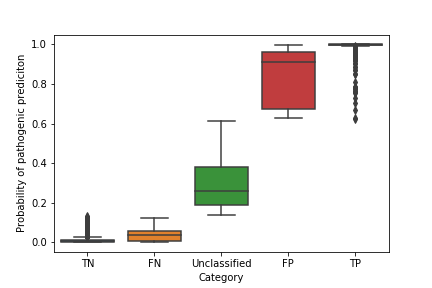

**Figure 7d.** Distribution of predicted duplication on testing data (Category) according to its probability of pathogenic prediction. (TN = True negative, FN = False negative, TP = True positive, FP = False positive)

**Tuning hyperparameters for XGBoost Classifier**

**Figure 8.** (Set of graphs) Tuning hyperparameters for XGBoost Classifier

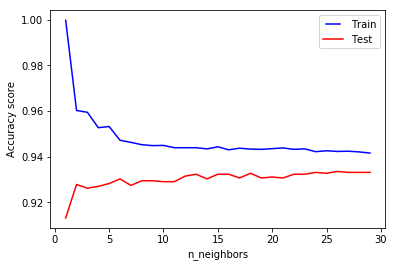

**Figure 8a.** Tuning number of neighbors according to accuracy score

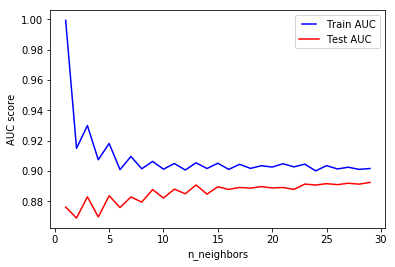

**Figure 8b.** Tuning number of neighbors according to AUC score.

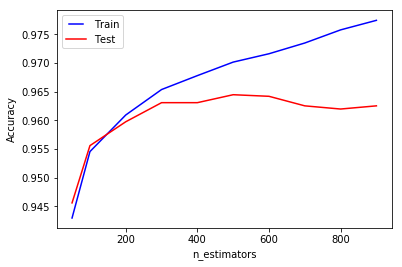

**Figure 8c.** Tuning number of estimators according to accuracy

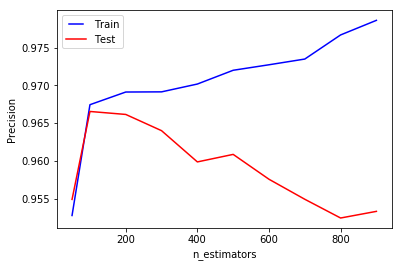

**Figure 8d.** Tuning number of estimators according to precision

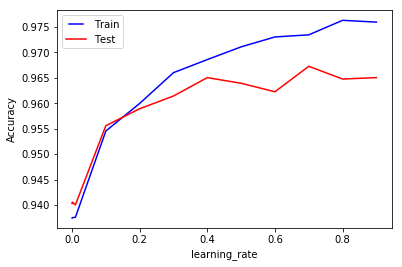

**Figure 8e.** Tuning number of learning rate according to accuracy

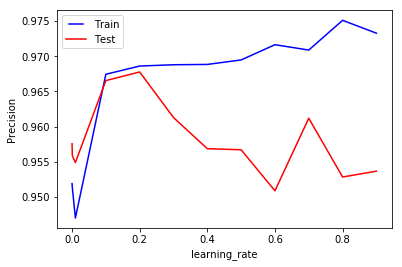

**Figure 8f.** Tuning number of learning rate according to precision

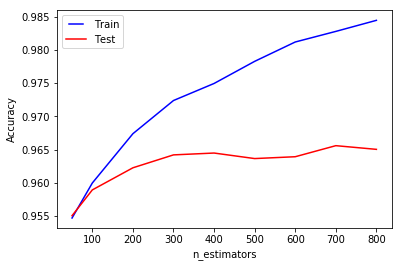

**Figure 8g.** Tuning number of estimators according to accuracy (learning rate set to 0.2)

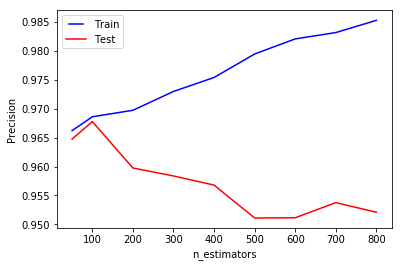

**Figure 8h.** Tuning number of estimators according to precision (learning rate = 0.2)

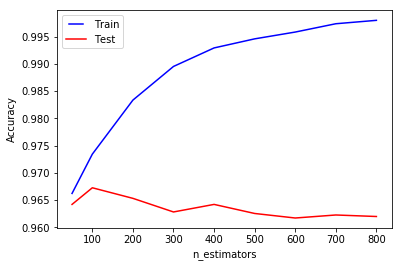

**Figure 8i.** Tuning number of estimators according to accuracy (learning rate = 0.7)

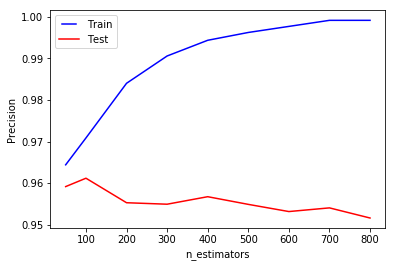

**Figure 8j.** Tuning number of estimators according to precision (learning rate = 0.7)

**Feature importances of used attributes**

| **Significance** | **Importance** | **Attribute** |
| --- | --- | --- |
| 1. | 0.33219743 | CG score |
| 2. | 0.2510013 | gene_count |
| 3. | 0.22639073 | morbidGene_count |
| 4. | 0.090607174 | EpLI score |
| 5. | 0.052542932 | DGV Frequency |
| 6. | 0.034897126 | HI score |
| 7. | 0.012363238 | EZ score |

**Table 4a**. Order of significance (feature importances) of used attributes for xgboost prediction model for deletions

| **Significance** | **Importance** | **Attribute** |
| --- | --- | --- |
| 1. | 0.64077955 | gene_count |
| 2. | 0.16930676 | EpLI score |
| 3. | 0.094157524 | morbidGene_count |
| 4. | 0.056282803 | DGV Frequency |
| 5. | 0.03947325 | EZ score |

**Table 4b.** Order of significance (feature importances) of used attributes for xgboost prediction model for duplications
